# Tumor Microenvironment in Ovarian Cancer through Spatial Transcriptomics and Identification of Key Gene Expression Profiles

**DOI:** 10.1101/2025.04.25.650590

**Authors:** Kevser Kübra Kırboğa, Ecir Uğur Küçüksille, Mithun Rudrapal, Emre Aktaş, Raghu Ram Achar, Gouri Deshpande, Victor Stupin, Ekaterina Silina

## Abstract

Ovarian cancer exhibits marked heterogeneity within its complex tumor microenvironment, driving progression, metastasis, and therapeutic resistance. This study presents a novel integration of spatial transcriptomics with single-cell RNA sequencing to comprehensively characterize the molecular landscape of ovarian cancer tissues. Employing the Leiden clustering algorithm at an optimal resolution of 0.7, we identified 13 distinct cellular subpopulations with unique transcriptional signatures. Pseudotime trajectory analysis revealed dynamic gene expression patterns associated with disease progression. Notably, FOSB, AMOTL2, and SLCO4A1 showed progressive upregulation during cellular differentiation, strongly implicating them in tumor cell motility and metastatic potential. Conversely, PLEK, CFB, and ADGRB1 maintained stable expression patterns, suggesting critical roles in maintaining cellular homeostasis and signal transduction. We observed significant downregulation of mitochondrial genes (particularly MT-CO2) and extracellular matrix components (such as COL1A2) in later differentiation stages, highlighting their importance in early oncogenic processes. Functional enrichment analysis connected these gene clusters to key biological pathways including extracellular matrix remodeling (COL family), mitochondrial energy metabolism (MT-CO2), and immune response modulation (CD163, PLEK). Our findings not only provide unprecedented insights into the molecular architecture of ovarian cancer but also identify several promising biomarkers and potential therapeutic targets. This integrative spatial-temporal approach establishes a robust framework for understanding the complex interplay within the tumor microenvironment and offers new strategic directions for developing personalized interventions in ovarian cancer treatment.

## 1. Introduction

Ovarian cancer is a complex and heterogeneous disease characterized by diverse cellular compositions and variable molecular profiles within the tumor microenvironment [1]. This heterogeneity significantly contributes to the pathophysiology of ovarian cancer, making the treatment and research of the disease particularly challenging [2]. The dynamic interactions between tumor cells and the surrounding stromal and immune cells play a crucial role in tumor progression, metastasis, and therapy resistance, further increasing the complexity of the disease [3]. Understanding the spatial organization of cells within the tumor microenvironment is crucial for elucidating the biological processes that support ovarian cancer progression [4, 5]. Spatial transcriptomics is an advanced technology that enables the mapping of gene expression profiles within tissue architecture. Unlike traditional bulk RNA sequencing techniques, which provide averaged gene expression profiles across a heterogeneous cell population, spatial transcriptomics preserves spatial context, allowing for the identification of gene expression patterns specific to certain regions or cell types within the tumor [6–8]. This approach is precious in diseases like ovarian cancer, where the spatial arrangements of cells and interactions between tumor, immune, and stromal cells play a critical role in disease progression and response to therapy [9, 10].

Recent advancements in spatial transcriptomics have revolutionized cancer research by facilitating gene expression analysis at the cellular level within tissues. This technology enhances our understanding of cellular heterogeneity within cancer tissues and the biological processes within the tumor microenvironment. For instance, Moncada et al. demonstrated the spatial architecture and heterogeneity of gene expression in pancreatic ductal adenocarcinomas (PDAC) [11]. Similarly, Lv et al. illustrated the utility of spatial transcriptomic analyses in invasive micropapillary carcinoma of the breast, revealing gene expression characteristics that underscore the tool’s effectiveness in examining the distribution and biological functions of cells within the tissue [12]. The heterogeneity of tumors has been identified as an important prognostic factor in breast and lung cancers [13], with spatial transcriptomics techniques revealing distinct gene expression profiles between the tumor core and their implications for tumor biology and patient survival [14]. Additionally, in non-small cell lung cancer (NSCLC), spatial transcriptomics has shed light on how macrophage distribution affects sensitivity and resistance to anti-PD1/PD-L1 antibodies [15].

Denisenko et al. highlighted that ovarian cancer subclones exhibit distinct gene expression profiles and cellular behaviors, with subclones creating unique microenvironments that support their growth and survival through autocrine signaling [9]. Stur and colleagues mapped the gene expression profiles across different high-grade serous ovarian carcinoma (HGSOC) tissue regions, examining the cellular organization within the tumor and the interactions between tumor subclones [10]. Similarly, Ju et al. integrate spatial transcriptomics and CT imaging to uncover molecular features of recurrent and non-recurrent HGSOC, providing a comprehensive view of the tumor microenvironment and its influence on disease progression. By correlating gene expression profiles with CT phenotypes, the study identifies critical pathways, such as TNF-α signaling via NF-κB and oxidative phosphorylation, that are linked to recurrence, while also pinpointing potential biomarkers for favorable prognosis, such as PTGDS, CXCL14, and RNASE1 [16]. Further extending this understanding, Yeh et al. applied single-cell spatial and perturbational transcriptomics to investigate immune evasion mechanisms in high-grade serous tubo-ovarian cancer [17]. Their study mapped the spatial organization of over 2.5 million cells from 130 tumors across 94 patients, revealing a malignant cell state influenced by tumor genetics that predicts T cell and natural killer cell infiltration levels and responsiveness to immune checkpoint blockade. By identifying genetic perturbations—such as PTPN1 and ACTR8 knockouts—that influence this malignant cell state, the study demonstrated how these alterations could sensitize ovarian cancer cells to cytotoxic immune responses. Adding to this growing body of research, Wang et al. performed a spatial transcriptomic analysis to investigate the earliest steps in HGSOC pathogenesis [18]. Their study focused on serous tubal intraepithelial carcinoma (STIC), a precursor lesion of HGSOC, comparing its transcriptomic profile with that of carcinoma and matched normal fallopian tube epithelium. They identified insulin-like growth factor binding protein-2 (IGFBP2) as a key gene reactivated during tumor development through DNA hypomethylation. IGFBP2 expression, regulated by epigenetic changes, was shown to be critical for the proliferation of tubal epithelial cells via the AKT pathway, particularly in postmenopausal, estrogen-deprived microenvironments. Knockdown experiments confirmed IGFBP2’s role in tumor initiation and progression.

Ovarian cancer has a highly heterogeneous structure, and understanding gene expression patterns in the tumor microenvironment is critical for disease progression and determining treatment targets. However, it is noteworthy that spatial transcriptomic analyses performed on ovarian tumor tissue are limited in the literature and have not been sufficiently integrated with advanced methods (e.g., gene clustering, functional enrichment, and pseudotime analyses). This deficiency prevents the full elucidation of dynamic processes in the ovarian tumor microenvironment and the genetic mechanisms affecting these processes. This study analyzes gene expression patterns in detail with spatial transcriptomic data, focusing only on ovarian tumor tissue. The study aims to fill this important research gap in the literature by identifying critical genes in the tumor tissue, correlating these genes with functional biological processes, and examining temporal dynamics (pseudotime). In this way, the aim is to understand better the cellular and molecular mechanisms in the ovarian tumor microenvironment.

## 2. Methods

### 2.1. Data Preparation and Filtering Process

This study utilized high-resolution spatial transcriptomic data from human ovarian cancer tissue obtained from 10x Genomics’ Xenium platform (accessed 10/08/2024). The datasets were fully anonymized before our access, with no identifying information available to researchers. As this study used only publicly available data without any direct human participant involvement, specific informed consent for our research was not required. The original tissue samples and data collection procedures were managed by 10x Genomics according to their ethical protocols [19].

To ensure robust biological inference, we implemented a systematic quality control and preprocessing pipeline following established protocols [19, 20]. Initial quality filtering removed suboptimal cellular profiles based on two primary criteria: cells expressing fewer than 200 genes were excluded to eliminate potentially degraded or poorly captured cells, and genes detected in fewer than 3 cells were removed to prevent spurious expression patterns from influencing downstream analyses.

To further refine our dataset, we evaluated mitochondrial gene content as a key quality metric, as excessive mitochondrial RNA often indicates cellular stress or degradation. Cells exhibiting mitochondrial gene ratios exceeding 5% were excluded from subsequent analyses. For the remaining high-quality cells, we performed normalization by scaling the total gene expression level of each cell to 10,000 counts, followed by logarithmic transformation to stabilize variance and approximate normal distribution—critical steps for ensuring valid statistical comparisons in downstream clustering and differential expression analyses.

### 2.2. Identification of Cell Cluster with Leiden Algorithm and PAGA-Based Relationship Analysis

To decipher the cellular heterogeneity within ovarian cancer tissues, we performed comprehensive clustering analysis on our spatial transcriptomic data. This approach enabled identification of distinct cellular subpopulations based on transcriptional similarities, providing critical insights into the complex cellular composition of the tumor microenvironment. We selected the Leiden algorithm for cell clustering due to its superior performance in handling large-scale single-cell datasets and its robust modularity optimization properties that produce biologically meaningful partitions [21]. To systematically determine the optimal clustering resolution, we performed a multi-resolution analysis at five distinct granularity levels (0.4, 0.6, 0.7, 0.8, and 1.0). This approach allowed us to evaluate the trade-off between under-clustering (at lower resolutions, which yields fewer but larger clusters) and over-clustering (at higher resolutions, which produces more detailed but potentially fragmented subgroups). Each resolution parameter setting was evaluated for biological coherence and statistical robustness.

The relationships between identified clusters were further characterized using Partition-based Graph Abstraction (PAGA), an advanced graph-based method that preserves both global and local topological structures within the data [22]. PAGA analysis generated a connectivity map between clusters, revealing potential differentiation trajectories and functional relationships that might not be apparent from clustering alone. The spatial organization and transcriptional relationships between cell clusters were visualized using PAGA-enhanced Uniform Manifold Approximation and Projection (UMAP) overlays [23], providing an integrated view of both cluster identities and their relative positions in the transcriptional landscape.

This integrated analytical framework—combining Leiden clustering with multi-resolution assessment and PAGA-based relationship mapping—significantly enhanced our ability to detect biologically meaningful cellular subpopulations and understand their functional interrelationships within the complex ovarian cancer ecosystem.

### 2.3. Differential Gene Expression Analysis

Following cluster identification, we conducted a comprehensive differential gene expression analysis across the 13 cell clusters defined at the optimal resolution of 0.7. This critical analytical step aimed to identify cluster-specific transcriptional signatures, characterize the underlying biological functions of each cellular subpopulation, and discover potential biomarkers with diagnostic or therapeutic relevance in ovarian cancer.

We employed the Wilcoxon rank-sum test [24], a robust non-parametric statistical method particularly suited for single-cell transcriptomic data as it does not assume normal distribution of gene expression values. For each gene across all clusters, we calculated multiple statistical metrics: log fold change (logFC) to quantify expression magnitude differences, raw p-values (pval) to assess statistical significance, and adjusted p-values (pval_adj) using the Benjamini-Hochberg method [25] o control for false discovery rate in multiple hypothesis testing.

The differential expression analysis was structured as a one-versus-all comparison, where gene expression in each cluster was compared against its expression in all other clusters combined. Using the Leiden-defined cluster assignments as the grouping variable, we systematically tested each gene for significant differential expression across all clusters. This methodical approach enabled us to identify gene sets uniquely upregulated or downregulated in specific cellular subpopulations, providing insights into cluster-specific biological functions and revealing potential cellular markers characteristic of each distinct cell type within the ovarian cancer microenvironment.

### 2.4. Functional Enrichment Analysis

To elucidate the biological significance of our identified cellular subpopulations, we performed a comprehensive functional enrichment analysis focusing on the most statistically significant genes from each cluster. For each of the 13 clusters, we selected the top five differentially expressed genes exhibiting the lowest p-values, representing the most reliable cluster-specific markers.

The analysis utilized GProfiler [26] a powerful functional profiling tool, to systematically interrogate these gene sets against the Homo sapiens reference genome. Our enrichment strategy incorporated an extensive collection of complementary biological knowledge databases to ensure comprehensive pathway and functional annotation. This multi-database approach included the three primary Gene Ontology categories (Biological Process [BP], Molecular Function [MF], and Cellular Component [CC] [27]), alongside established pathway resources including KEGG [28], and Reactome [29],Additionally, the analysis incorporated specialized databases such as Human Phenotype Ontology (HPO), DisGeNET [30], for disease associations, miRTarBase [31], for microRNA interactions, WikiPathways [32], for community-curated pathways, and protein interaction resources CORUM [33], and BioGRID [34], This integrative enrichment framework allowed for characterization of each cluster’s functional profile across multiple biological dimensions simultaneously: cellular processes, molecular activities, subcellular localizations, disease associations, metabolic pathways, regulatory networks, and protein interaction complexes. Statistically significant enrichment terms were identified using adjusted p-values to control for multiple testing effects.

The most significant functional enrichment results for each cluster are presented in **Table 2**, revealing distinct biological signatures that define each cellular subpopulation within the ovarian cancer microenvironment. Complete enrichment results for all analyzed genes are provided in Supplementary **Table S2**. This multi-dimensional functional characterization provided crucial insights into the specialized biological roles of each cellular subtype identified in our spatial transcriptomic analysis.

### 2.5. Pseudotime Analysis to Cellular Differentiation Processes

To gain insight into the temporal dynamics of cellular differentiation and biological pathway progression in ovarian cancer, pseudotime analysis was implemented on a curated set of biologically significant genes [35]. The analysis focused on 13 key genes (OLIG3, FOSB, AMOTL2, SLCO4A1, MT-CO2, EPCAM, PLEK, CFB, COL1A2, ADGRB1, CD163, ARHGEF1, VWF) selected based on their statistical significance (high logFC values and low p-values) across the identified clusters.

The analytical pipeline began with data pre-processing, including normalization, logarithmic transformation, and scaling of gene expression values to ensure comparability across cells. Cellular relationships were then modeled through construction of a neighborhood graph, followed by application of the Leiden clustering algorithm at multiple resolution parameters (0.4, 0.6, 0.7, 0.8, and 1.0) to identify the optimal clustering granularity that best captured biologically meaningful cellular states.

Inter-cluster transitions and differentiation trajectories were subsequently mapped using Partition-based Graph Abstraction (PAGA) analysis. The root cell for pseudotime ordering was carefully selected based on three key criteria: biological context relevance, connection density within the PAGA graph, and centrality among graph nodes—ensuring that developmental trajectories were anchored at an appropriate starting point.

The Diffusion Pseudotime (DPT) algorithm [36] was applied to calculate developmental sequences, ordering cells along continuous trajectories starting from the designated root cell. This approach enabled reconstruction of potential differentiation paths based on transcriptional similarities, revealing how cells might progress through various states during tumor development.

To thoroughly investigate transition dynamics, cells falling within specific pseudotime intervals were defined as “transition cells” and subjected to detailed expression profiling. Correlation analysis between gene expression patterns across these transition cells was visualized through heatmaps, revealing coordinated expression changes during cellular differentiation. Additionally, expression trajectories for each of the 13 selected genes were plotted against pseudotime progression, allowing identification of genes with consistent upregulation (e.g., FOSB, AMOTL2), stable expression (e.g., PLEK, CFB), or progressive downregulation (e.g., MT-CO2, COL1A2) during the differentiation process.

This temporal reconstruction of gene expression dynamics provided valuable insights into the molecular mechanisms underlying cellular state transitions in the ovarian cancer microenvironment.

## 3. Results

### 3.1. Data Quality Control and Preprocessing

Our analysis began with a comprehensive spatial transcriptomic dataset comprising 6,566 cells and 18,082 genes derived from human ovarian cancer tissue. To ensure robust and reliable biological inference, we implemented a stringent quality control and preprocessing pipeline with multiple sequential filtering steps.

Initial quality filtration removed cells expressing fewer than 200 unique genes, as these likely represented low-quality captures or cellular debris. Simultaneously, genes detected in fewer than 3 cells were excluded to eliminate potential technical artifacts and spurious expression signals. This preliminary filtering retained 6,561 cells (99.9% of initial cells) and 16,753 genes (92.6% of initial genes).

Further quality assessment evaluated mitochondrial gene content—an established marker of cell stress and technical artifacts in single-cell analyses. Cells exhibiting mitochondrial content exceeding 5% were excluded, resulting in a high-quality dataset of 6,264 cells (95.4% of original cells) with consistent technical characteristics.

The normalized expression data underwent logarithmic transformation to stabilize variance across expression levels and approximate normal distribution for downstream statistical analyses. Feature selection then identified 2,773 highly variable genes that maximally captured biological variation between cells, substantially reducing dimensionality while preserving biologically meaningful signals.

Regression analysis removed technical confounders, including total gene counts and mitochondrial content, minimizing their impact on gene expression patterns. Final scaling standardized expression values to unit variance, with upper thresholds set at 10 to prevent outlier effects on subsequent analyses.

This systematic preprocessing approach yielded a high-fidelity dataset optimized for detailed characterization of cellular heterogeneity and gene expression dynamics within the ovarian cancer microenvironment, establishing a solid foundation for downstream clustering and trajectory analyses.

### 3.2. Identification of Cell Subpopulations and PAGA-Based Relationship Analysis

Our comparative multi-resolution clustering analysis (Fig. 1) revealed Resolution 0.7 as the optimal parameter for characterizing cellular heterogeneity within the ovarian cancer microenvironment. This systematic evaluation across different granularity levels (0.4, 0.6, 0.7, 0.8, and 1.0) demonstrated a critical trade-off between under-clustering and over-fragmentation of cellular populations.

**Fig. 1.**
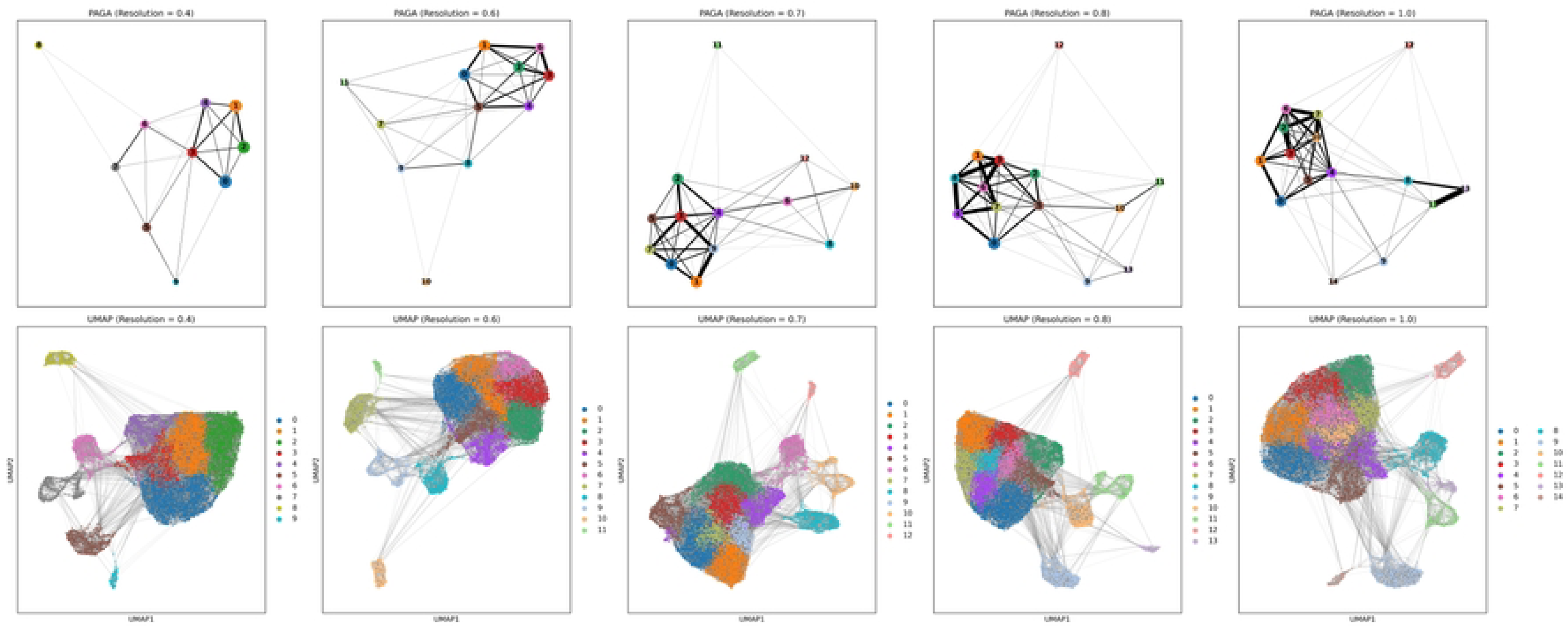
Comparison of PAGA and UMAP Analyses Across Different.

At the lower Resolution 0.4, the algorithm identified only 10 broad cellular groupings. While these clusters captured major cell type divisions, they lacked sufficient granularity to distinguish important biological subpopulations known to exist in ovarian cancer tissues. This coarse clustering merged transcriptionally distinct cell states, potentially obscuring functionally relevant cellular subtypes and their specific contributions to tumor biology.

Conversely, higher Resolutions 0.8 and 1.0 produced excessive fragmentation, generating 12-14 clusters including several small groups (containing <50 cells) with questionable biological significance. These higher resolutions introduced cluster boundaries that did not reflect meaningful biological distinctions, as evidenced by their poor connectivity patterns in the PAGA topology graph. The resulting over-partitioning created artificial separations between transcriptionally similar cells, complicating biological interpretation.

Resolution 0.7 emerged as the biologically optimal parameter, yielding 13 well-defined clusters with clear transcriptional boundaries while maintaining biologically plausible interconnections. PAGA visualization at this resolution revealed coherent trajectories and hierarchical relationships between clusters that aligned with expected biological transitions in tumor tissue. Each cluster contained sufficient cells for robust statistical analysis while preserving distinct transcriptional identities.

The connectivity patterns at Resolution 0.7 displayed strong correspondence with known biological relationships between cell types in the tumor microenvironment, enabling confident interpretation of cellular hierarchies and potential differentiation trajectories. This resolution provided the ideal balance between analytical resolution and biological coherence, establishing a solid foundation for downstream analyses of differential gene expression and functional characterization.

### 3.3. Differential Gene Expression Analysis

The differential gene expression analysis across the 13 cellular clusters (Resolution 0.7) revealed distinctive transcriptional signatures characterizing each subpopulation within the ovarian cancer microenvironment (**Table 1**). Each cluster exhibited a unique gene expression profile with remarkably high statistical significance, suggesting robust biological distinctions between the identified cell populations. Cluster 0 was primarily defined by the pronounced expression of neural-lineage regulator OLIG3 (LogFC: 3.001; p-value: 1.8E-108) and vesicular trafficking mediator RAB2A (LogFC: 2.971; p-value: 8.23E-96), suggesting potential neurodevelopmental characteristics within this cellular subpopulation. Stress-responsive transcription factor FOSB (LogFC: 3.028; p-value: 5.7E-179) and growth differentiation factor GDF15 (LogFC: 2.741; p-value: 7.4E-165) dominated the transcriptional landscape of Cluster 1, indicating potential roles in stress adaptation and growth regulation pathways. Cluster 2 demonstrated significant upregulation of cell junction regulator AMOTL2 (LogFC: 2.833; p-value: 3.4E-180) and extracellular matrix component LAMC2 (LogFC: 3.102; p-value: 7.2E-168), suggesting involvement in cell adhesion and epithelial organization.

**Table 1.** Differential Gene Expression Results for Clusters Identified by Leiden Algorithm (Full List Provided in Supplementary Table S1).

| Cluster | Gene | LogFC | pval | pval_adj |
| --- | --- | --- | --- | --- |
| 0 | OLIG3 | 3.001352 | 1.8E-108 | 4.9E-105 |
| 0 | RAB2A | 2.971275 | 8.23E-96 | 1.12E-92 |
| 0 | AC007906.2 | 2.008865 | 1.71E-94 | 1.55E-91 |
| 0 | ZC3HAV1 | 1.90045 | 2.13E-88 | 1.45E-85 |
| 0 | CADM1 | 1.973429 | 2.2E-82 | 1.2E-79 |
| 1 | FOSB | 3.028073 | 5.7E-179 | 1.5E-175 |
| 1 | GDF15 | 2.740653 | 7.4E-165 | 1E-161 |
| 1 | HMGCR | 2.536022 | 6.4E-151 | 5.8E-148 |
| 1 | VTCN1 | 2.553989 | 8.3E-137 | 5.6E-134 |
| 1 | TXNIP | 2.205701 | 1.2E-129 | 6.8E-127 |
| 2 | AMOTL2 | 2.833404 | 3.4E-180 | 9.1E-177 |
| 2 | ITGA5 | 2.765204 | 2E-174 | 2.7E-171 |
| 2 | LAMC2 | 3.10224 | 7.2E-168 | 6.5E-165 |
| 2 | SLCO4A1 | 2.464803 | 1.4E-156 | 9.4E-154 |
| 2 | LAMB3 | 2.766597 | 1E-155 | 5.7E-153 |
| 3 | SLCO4A1 | 3.072567 | 4.9E-204 | 1.3E-200 |
| 3 | PADI1 | 3.85658 | 8E-166 | 1.1E-162 |
| 3 | ANGPTL4 | 2.819982 | 1.3E-150 | 1.1E-147 |
| 3 | SERPINB9 | 2.705786 | 9.8E-140 | 6.7E-137 |
| 3 | ITPKC | 2.482106 | 7.7E-131 | 4.2E-128 |
| 4 | MT-CO2 | 3.024452 | 5.25E-80 | 1.43E-76 |
| 4 | MT-CO3 | 3.037517 | 8.59E-73 | 1.17E-69 |
| 4 | MT-ND4 | 2.800635 | 8.02E-69 | 7.27E-66 |
| 4 | MT-ATP6 | 2.966777 | 3.11E-64 | 2.11E-61 |
| 4 | MT-CYB | 3.112972 | 6.51E-64 | 3.54E-61 |
| 5 | EPCAM | 3.501746 | 2.7E-197 | 7.5E-194 |
| 5 | CST3 | 3.619715 | 3.2E-186 | 4.3E-183 |
| 5 | LAMP1 | 3.269331 | 1E-170 | 9.2E-168 |
| 5 | COL4A1 | 2.899471 | 3.6E-165 | 2.4E-162 |
| 5 | CD63 | 2.984298 | 1.9E-163 | 1E-160 |
| 6 | PLEK | 7.273333 | 4.4E-176 | 1.2E-172 |
| 6 | MXD1 | 5.558567 | 2.5E-168 | 3.4E-165 |
| 6 | ITGAX | 6.134906 | 1E-160 | 9.1E-158 |
| 6 | SQSTM1 | 3.604392 | 1.8E-160 | 1.2E-157 |
| 6 | PLAUR | 4.806293 | 8.4E-155 | 4.6E-152 |
| 7 | CFB | 3.285925 | 1.2E-99 | 3.29E-96 |
| 7 | BIRC3 | 2.676786 | 3.77E-83 | 5.13E-80 |
| 7 | ATP6V1C2 | 2.091236 | 2.14E-60 | 1.94E-57 |
| 7 | NAMPT | 1.585726 | 1.57E-37 | 1.07E-34 |
| 7 | LEFTY1 | 1.750327 | 5.23E-37 | 2.85E-34 |
| 8 | COL1A2 | 8.016911 | 2.7E-216 | 7.4E-213 |
| 8 | COL3A1 | 8.52774 | 3.6E-215 | 4.9E-212 |
| 8 | COL6A2 | 6.640284 | 2E-213 | 1.8E-210 |
| 8 | COL1A1 | 7.077699 | 1.4E-205 | 9.7E-203 |
| 8 | COL6A3 | 8.566475 | 2.2E-190 | 1.2E-187 |
| 9 | ADGRB1 | 3.194814 | 2.35E-91 | 6.38E-88 |
| 9 | NID2 | 2.464078 | 6.48E-82 | 8.81E-79 |
| 9 | CCDC91 | 2.365793 | 1.16E-63 | 1.05E-60 |
| 9 | LOX | 2.155361 | 3.98E-59 | 2.71E-56 |
| 9 | KISS1R | 2.606966 | 8.01E-58 | 4.36E-55 |
| 10 | CD163 | 9.192554 | 8.4E-160 | 2.3E-156 |
| 10 | SRGN | 6.185234 | 2.9E-148 | 3.9E-145 |
| 10 | STAB1 | 8.639015 | 3.4E-135 | 3.1E-132 |
| 10 | HK3 | 7.186135 | 3E-130 | 2E-127 |
| 10 | LILRB1 | 7.763309 | 5.5E-128 | 3E-125 |
| 11 | ARHGEF1 | 4.041759 | 3.99E-78 | 1.08E-74 |
| 11 | AKNA | 4.561586 | 3.08E-77 | 4.19E-74 |
| 11 | ACAP1 | 7.593784 | 1.08E-75 | 9.78E-73 |
| 11 | TMC8 | 7.167469 | 3.5E-72 | 2.38E-69 |
| 11 | IKZF1 | 7.272673 | 1.36E-64 | 7.41E-62 |
| 12 | VWF | 10.78681 | 5.82E-49 | 1.58E-45 |
| 12 | EGFL7 | 8.457527 | 4.9E-46 | 6.66E-43 |
| 12 | SPARC | 7.585188 | 5.62E-45 | 5.09E-42 |
| 12 | CD93 | 7.667711 | 2.02E-41 | 1.38E-38 |
| 12 | TMEM255B | 8.529364 | 3.69E-41 | 2.01E-38 |

The strongest differential expression was observed in Cluster 3, marked by PADI1 (LogFC: 3.857; p-value: 8.0E-166) and membrane transporter SLCO4A1 (LogFC: 3.073; p-value: 4.9E-204), potentially indicating specialized metabolic functions within this subpopulation. Cluster 4 exhibited a distinctive signature of mitochondrial genes, including MT-CO2 (LogFC: 3.024; p-value: 5.25E-80) and MT-CYB (LogFC: 3.113; p-value: 6.51E-64), highlighting a potential metabolic specialization associated with oxidative phosphorylation. Epithelial cell marker EPCAM (LogFC: 3.502; p-value: 2.7E-197) and lysosomal protease inhibitor CST3 (LogFC: 3.620; p-value: 3.2E-186) characterized Cluster 5, suggesting an epithelial-derived subpopulation with specialized endolysosomal activity. Cluster 6 showed significant enrichment of immune-related genes, including ITGAX (LogFC: 6.135; p-value: 1.0E-160) and autophagy regulator SQSTM1 (LogFC: 3.604; p-value: 1.8E-160), pointing to a myeloid-lineage subpopulation. Complement factor CFB (LogFC: 3.286; p-value: 1.2E-99) dominated Cluster 7, suggesting involvement in innate immune response pathways. Cluster 8 was markedly characterized by extraordinarily high expression of multiple collagen family members, including COL1A2 (LogFC: 8.017; p-value: 2.7E-216) and COL3A1 (LogFC: 8.528; p-value: 3.6E-215), strongly indicating a fibroblast or stromal cell population. The cellular adhesion receptor ADGRB1 (LogFC: 3.195; p-value: 2.35E-91) and extracellular matrix modulator LOX (LogFC: 2.155; p-value: 3.98E-59) defined Cluster 9, suggesting roles in matrix remodeling. Cluster 10 exhibited extreme upregulation of macrophage marker CD163 (LogFC: 9.193; p-value: 8.4E-160) and immunoregulatory receptor LILRB1 (LogFC: 7.763; p-value: 5.5E-128), clearly identifying a tumor-associated macrophage population. Lymphocyte lineage transcription factor IKZF1 (LogFC: 7.273; p-value: 1.36E-64) and signaling regulator ACAP1 (LogFC: 7.594; p-value: 1.08E-75) were hallmarks of Cluster 11, indicating a lymphoid-derived population.

Finally, Cluster 12 was distinctively characterized by extreme expression of endothelial markers VWF (LogFC: 10.787; p-value: 5.82E-49) and EGFL7 (LogFC: 8.458; p-value: 4.9E-46), definitively identifying an endothelial cell population within the tumor microenvironment.

These cluster-specific gene signatures provide precise molecular definitions for each cellular subpopulation, enabling functional annotation and biological interpretation of the heterogeneous cellular composition within ovarian cancer tissues.

### 3.4. Functional Enrichment Analysis

Comprehensive functional enrichment analysis of the top differentially expressed genes from each cluster revealed distinct biological signatures and specialized cellular functions within the ovarian cancer ecosystem (**Table 2**, complete results in **Supplementary Table S2**). This analysis provided crucial insights into the functional architecture of the tumor microenvironment by linking transcriptional profiles to specific biological pathways and processes. Cluster 0 showed significant enrichment for POU3F1 transcription factor binding motifs (p-value: 0.012), suggesting potential neurodevelopmental regulatory programs active in this cellular subpopulation. This enrichment aligns with the high expression of OLIG3, indicating potential neural lineage-like characteristics within these tumor cells.

**Table 2.**
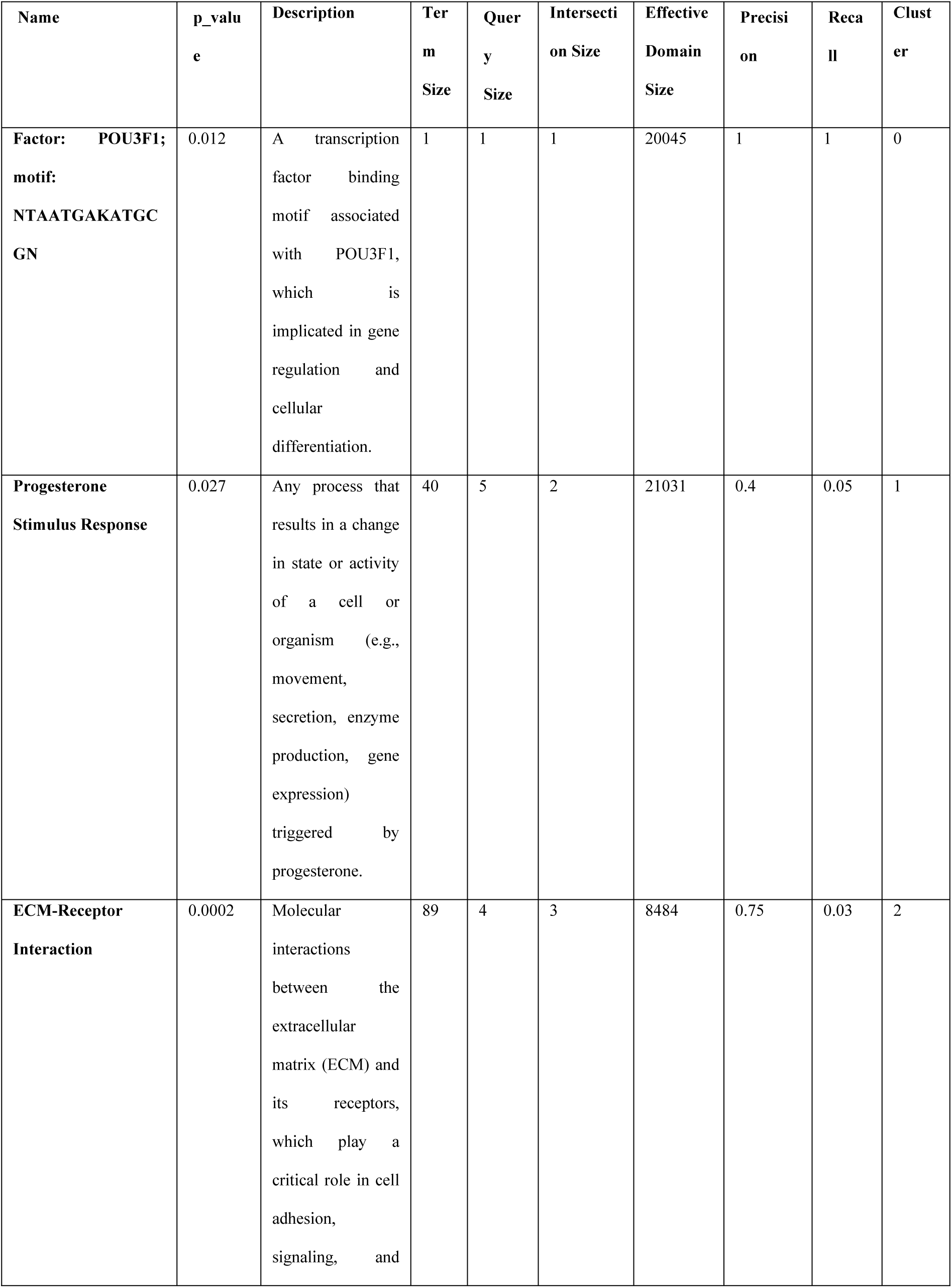
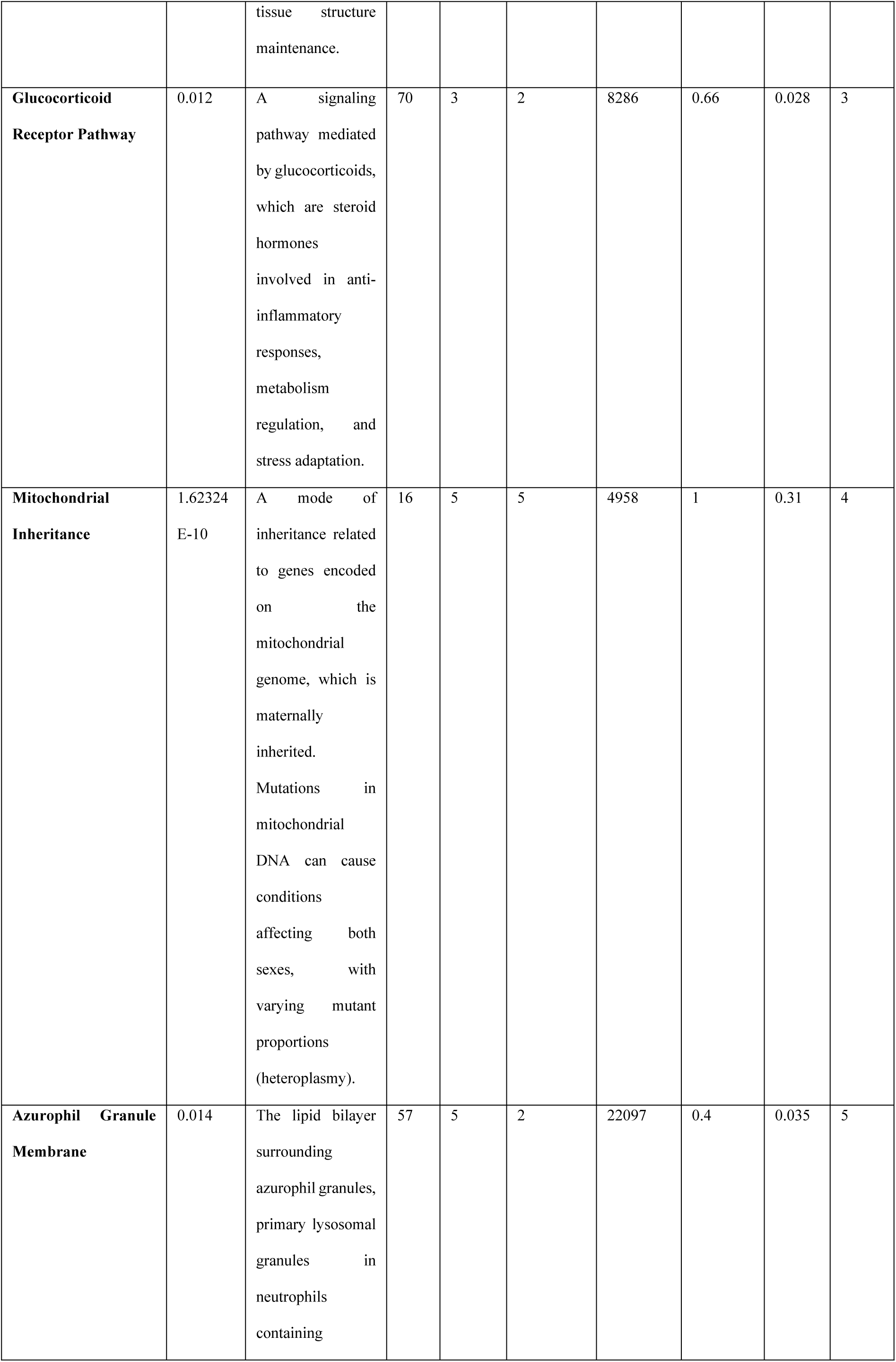
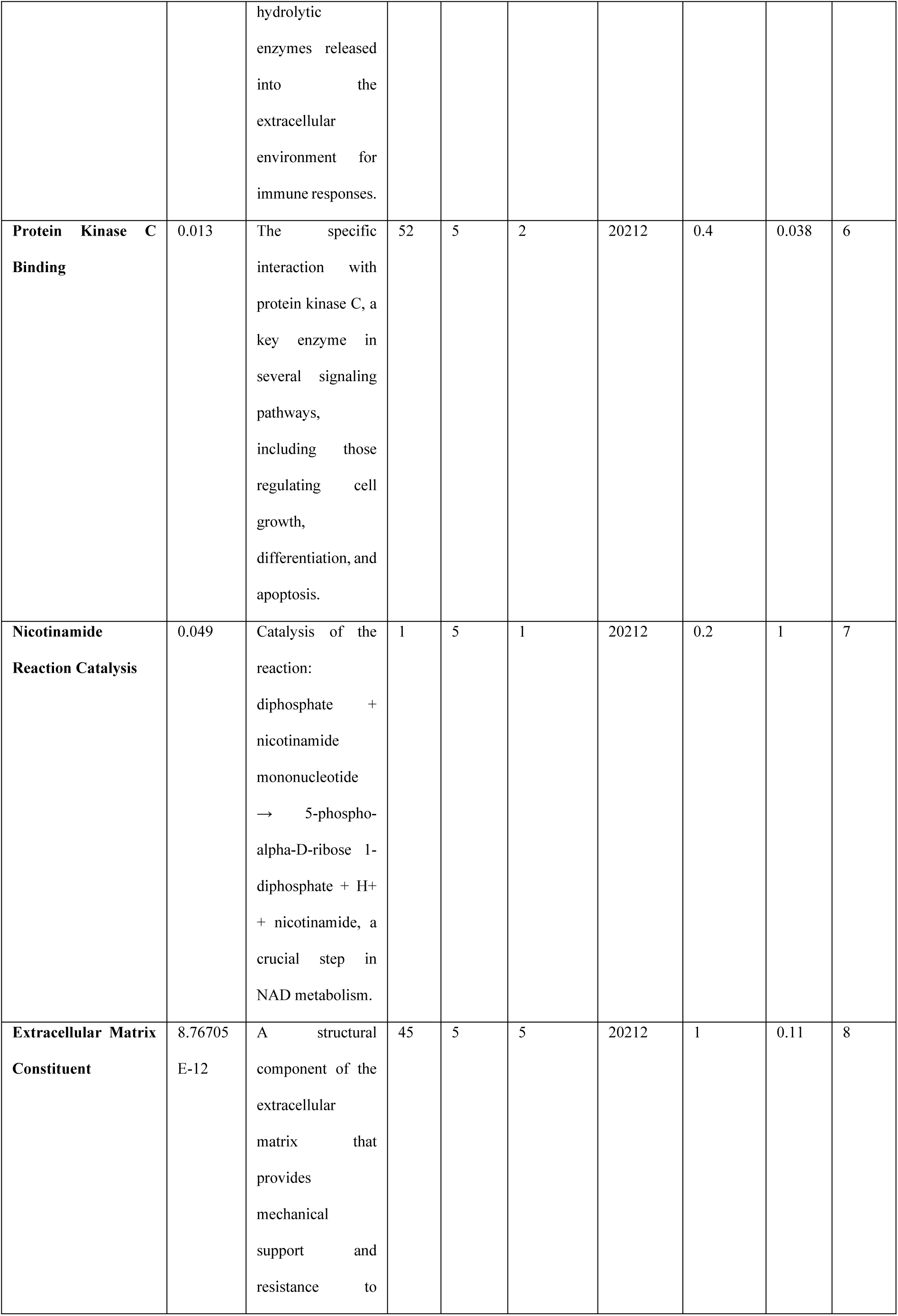
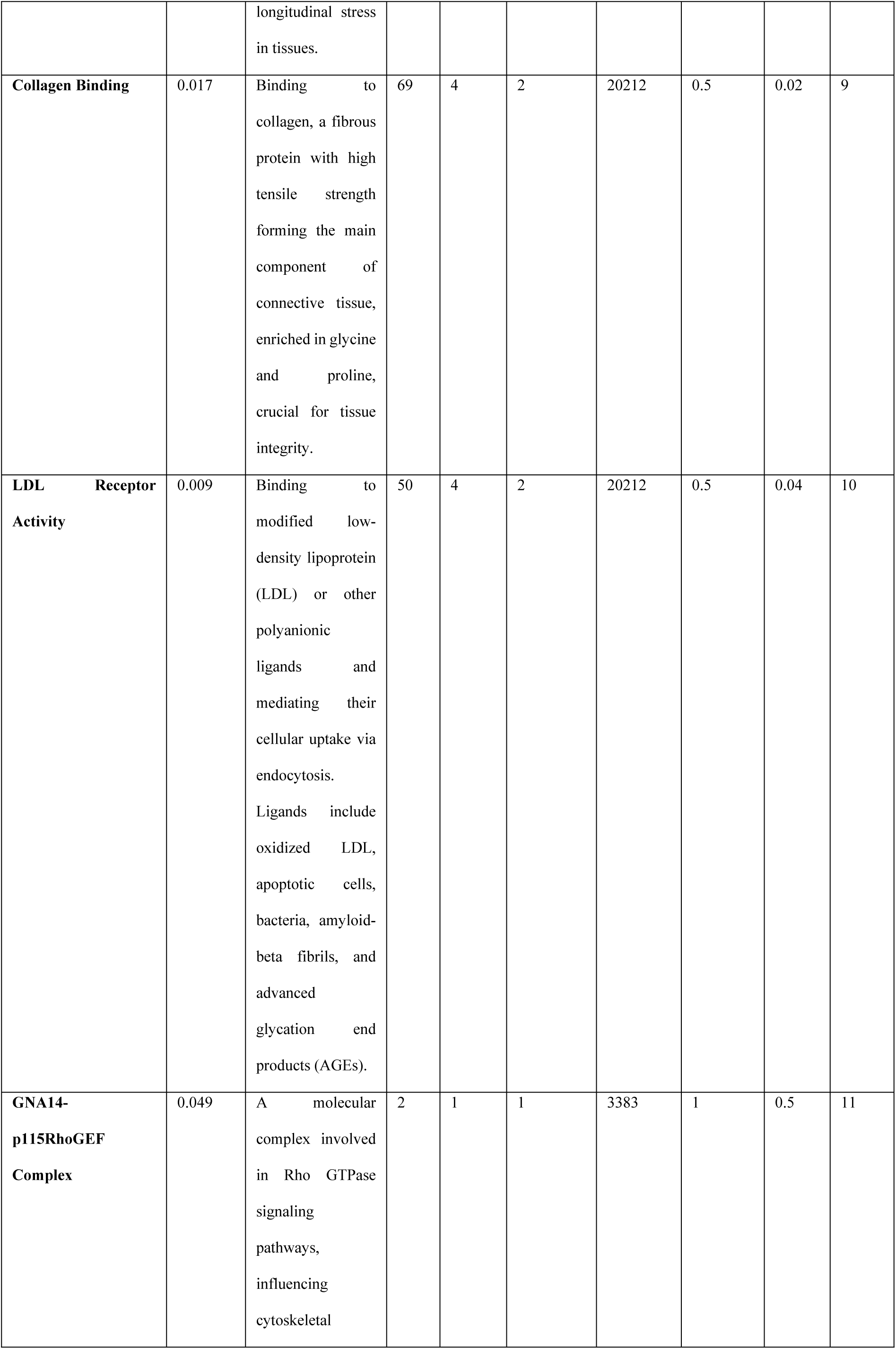
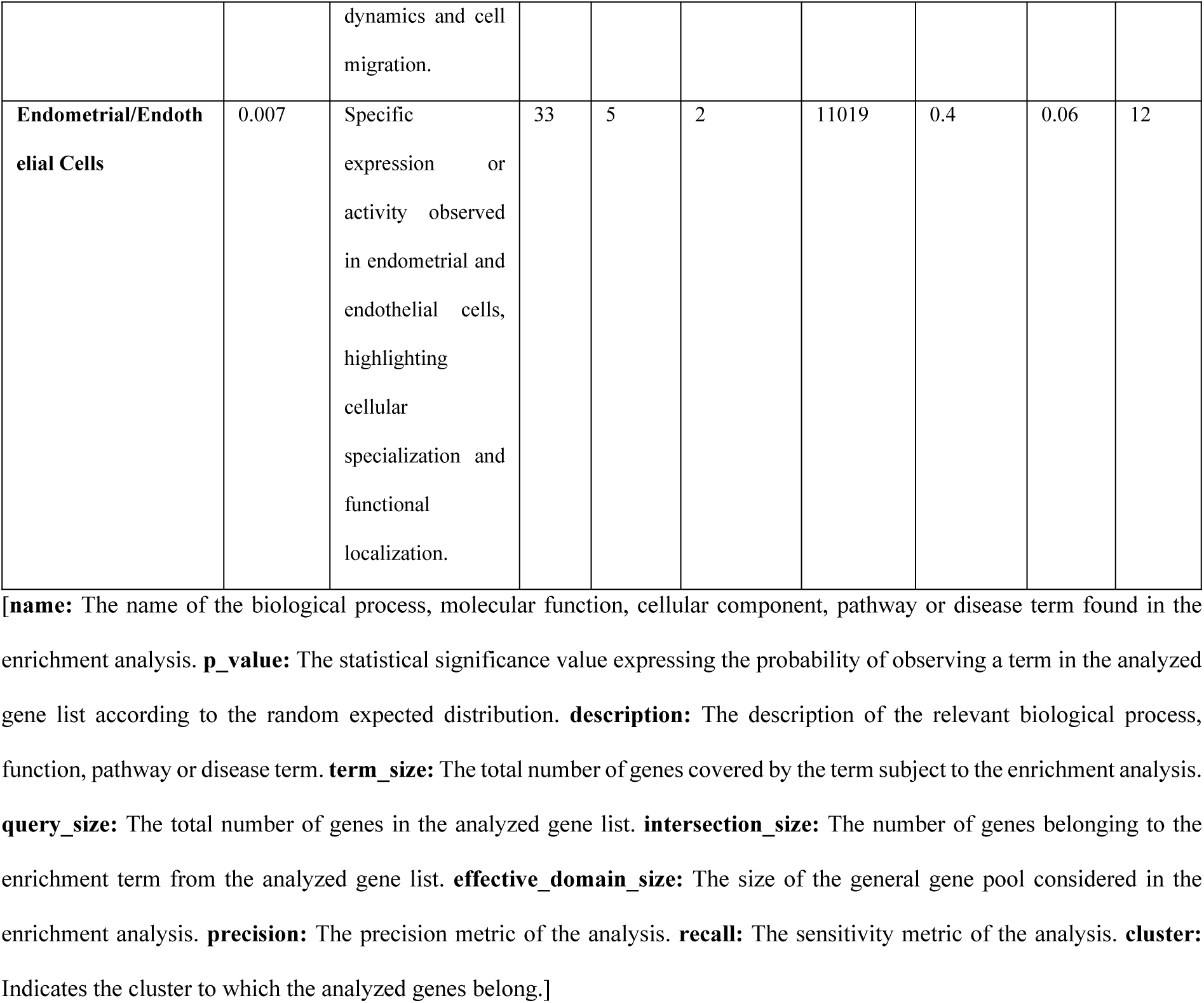
Functional Enrichment Analysis of Top 5 Genes from Each Cluster. The remaining analysis results are presented in Supplementary Table S2.

Hormone responsiveness emerged as the defining functional characteristic of Cluster 1, with significant enrichment for progesterone stimulus response pathways (p-value: 0.027). This hormonal sensitivity correlates with the high expression of stress-responsive transcription factor FOSB, potentially indicating an adaptive stress response mechanism operating in these cells.

Cluster 2 exhibited strong enrichment for ECM-receptor interaction pathways (p-value: 0.0002), highlighting the importance of cell-matrix communications in this subpopulation. This finding, coupled with high AMOTL2 and LAMC2 expression, suggests a cellular identity associated with tissue organization and structural integrity maintenance. The glucocorticoid receptor pathway was significantly enriched in Cluster 3 (p-value: 0.012), indicating specialized roles in anti-inflammatory processes and metabolic regulation. This pathway enrichment provides functional context for the high

SLCO4A1 expression observed in this cluster. Remarkably strong enrichment for mitochondrial inheritance patterns (p-value: 1.62E-10) characterized Cluster 4, perfectly aligning with its distinctive expression of multiple mitochondrial genes (MT-CO2, MT-CYB). This finding suggests a subpopulation with specialized metabolic functions dependent on mitochondrial activity. Cluster 5 showed significant enrichment for azurophil granule membrane components (p-value: 0.014), linking this subpopulation to specialized lysosomal functions involved in immune response. This enrichment complements the high expression of lysosomal protease inhibitor CST3 observed in this cluster. Protein kinase C binding functions were significantly enriched in Cluster 6 (p-value: 0.013), suggesting active roles in signal transduction pathways. Additionally, this cluster showed enrichment for nicotinamide reaction catalysis (p-value: 0.049), indicating specialized metabolic functions. Collagen binding emerged as the key functional attribute of Cluster 7 (p-value: 0.017), suggesting involvement in tissue organization and extracellular matrix interactions. This function complements the role of highly expressed complement factor CFB in this cluster. Cluster 8 demonstrated extremely significant enrichment for extracellular matrix constituent functions (p-value: 8.77E-12), perfectly aligning with its extraordinarily high expression of multiple collagen family members and confirming its identity as a stromal/fibroblast population. LDL receptor activity and associated endocytosis processes were significantly enriched in Cluster 9 (p-value: 0.009), suggesting specialized roles in lipid metabolism and cellular uptake mechanisms potentially relevant to tumor metabolism. Cluster 10 showed significant enrichment for GNA14-p115RhoGEF complex components (p-value: 0.049), implicating Rho GTPase signaling pathways in this macrophage-like subpopulation identified by high CD163 expression. Mechanisms regulating cell mobility and cytoskeletal dynamics were prominent in Cluster 11, supporting its potential identity as a lymphoid-derived population with significant migration capabilities.

Endometrial and endothelial cell-specific functions were significantly enriched in Cluster 12 (p-value: 0.007), perfectly aligning with its high expression of endothelial markers VWF and EGFL7 and confirming its identity as an endothelial subpopulation within the tumor micro-environment.

### 3.5. Pseudotime Analysis to Cellular Differentiation Processes

The expression profiles of 13 selected genes (OLIG3, FOSB, AMOTL2, SLCO4A1, MT-CO2, EPCAM, PLEK, CFB, COL1A2, ADGRB1, CD163, ARHGEF1, VWF) were examined during cellular differentiation and pseudotime. PAGA and UMAP images at each resolution (0.4, 0.6, 0.7, 0.8, 1.0) showed that cell clusters were organized differently. At resolution 0.4, 3 clusters were identified and a simple connectivity graph was observed throughout pseudotime. As the resolution increased (e.g., 0.7 and 1.0), the number of clusters increased, and the connections became more complex. In particular, the structure observed at resolution 0.7 presented a balanced view between biological significance and clustering and was considered an ideal result where transitional cells were clearly resolved. In pseudotime images, cells were sorted along the differentiation and the distribution of transitional cells (0.4–0.6 range) was resolved. The intensity of gene expressions for transitional cells was visualized at each resolution. At resolution 0.4, COL1A2, MT-CO2 and VWF genes showed significantly higher expression in some clusters, suggesting that these genes may play critical roles in differentiation processes. At resolutions 0.6 and 0.7, gene expression patterns were resolved in more detail, revealing distinct genetic profiles of transitional cells. It was determined that the intense expression of COL1A2, especially at resolution 0.7, may indicate the transition of cells in a specific differentiation pathway. At higher resolutions (0.8 and 1.0), cell clusters and gene expression patterns became more complex, and expressions of other genes such as EPCAM and FOSB were observed to be prominent in certain clusters. These analyses allowed the identification of different genetic features for transitional cells at each resolution. The changes of 13 genes during pseudotime were presented in a line graph, and these graphs detailed the fluctuations and common patterns in the expression patterns of the genes. For example, FOSB and AMOTL2 genes showed a significant increase during pseudotime, suggesting that these genes may have a potential role in cell differentiation and transition processes. In contrast, genes such as PLEK, CFB and CD163 generally exhibited a stable expression pattern and were observed to be less effective during differentiation. Genes such as COL1A2 and VWF were intensely expressed in specific transitional cells, suggesting that these genes may play critical roles at certain points in the differentiation process. In addition, it was observed that genes associated with mitochondrial and epithelial differentiation, such as MT-CO2 and EPCAM, generally fluctuated during pseudotime and may be active during processes where cells tend to a certain subset. Correlation analysis provided a comprehensive picture of the relationships between gene expressions. High positive correlations such as 0.52 between OLIG3 and AMOTL2 and 0.37 between OLIG3 and ARHGEF1 suggest that these genes may be involved in common biological processes. Similarly, positive correlations between EPCAM and MT-CO2 (0.34), CFB and ARHGEF1 (0.34) suggest that these genes are associated with common pathways or regulatory mechanisms. On the other hand, negative correlations are notably clustered around VWF; for example, values such as −0.44 between MT-CO2 and VWF and −0.34 between AMOTL2 and VWF suggest that these genes are involved in biologically opposite processes. The negative correlation of −0.30 between PLEK and SLCO4A1 suggests that these genes may play opposing roles in cellular differentiation. In contrast, the weak correlations observed between FOSB and other genes (e.g., −0.02 between FOSB and PLEK, 0.01 between FOSB and EPCAM) suggest that these genes participate in independent processes. The overall weak correlations between COL1A2 and other genes suggest that this gene may serve a specific function. Overall, this analysis suggests that genes such as OLIG3, AMOTL2, ARHGEF1, and EPCAM may play coordinated roles in common processes, whereas genes such as VWF may exhibit different or independent effects (**Fig. 2**).

**Fig. 2.**
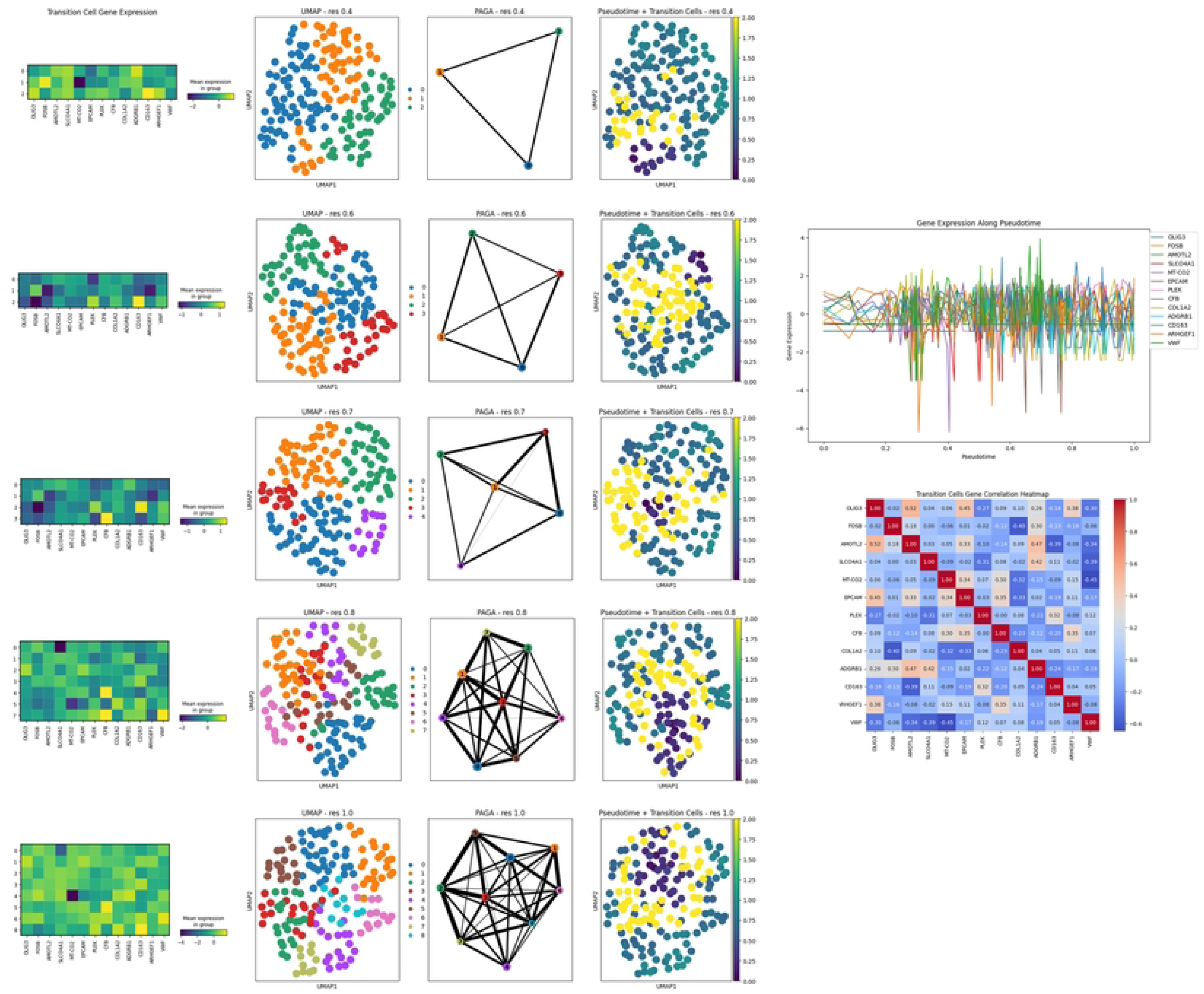
Comprehensive Analysis of Pseudotime Trajectories, Clust.

In **Fig. 3**, the expression distribution of each gene on UMAP is shown. OLIG3 shows low to moderate expression in a wide area, while FOSB shows high expression in a more limited area. AMOTL2 shows intense expression in multiple areas between clusters, while SLCO4A1 has a lower and more localized expression profile. MT-CO2 stands out with its distinctly high expression and shows activity in a wide cell population. EPCAM has distinct and intense expression regions in specific cell groups. PLEK and CFB are concentrated in more limited and localized regions. COL1A2 shows high expression peaks, especially in one or two areas. ADGRB1 shows moderate to high expression in multiple localized regions. CD163 is notable with high expression areas, while ARHGEF1 is concentrated in a smaller group of cells. Finally, VWF stands out with intense expression regions localized in specific clusters.

**Fig. 3.**
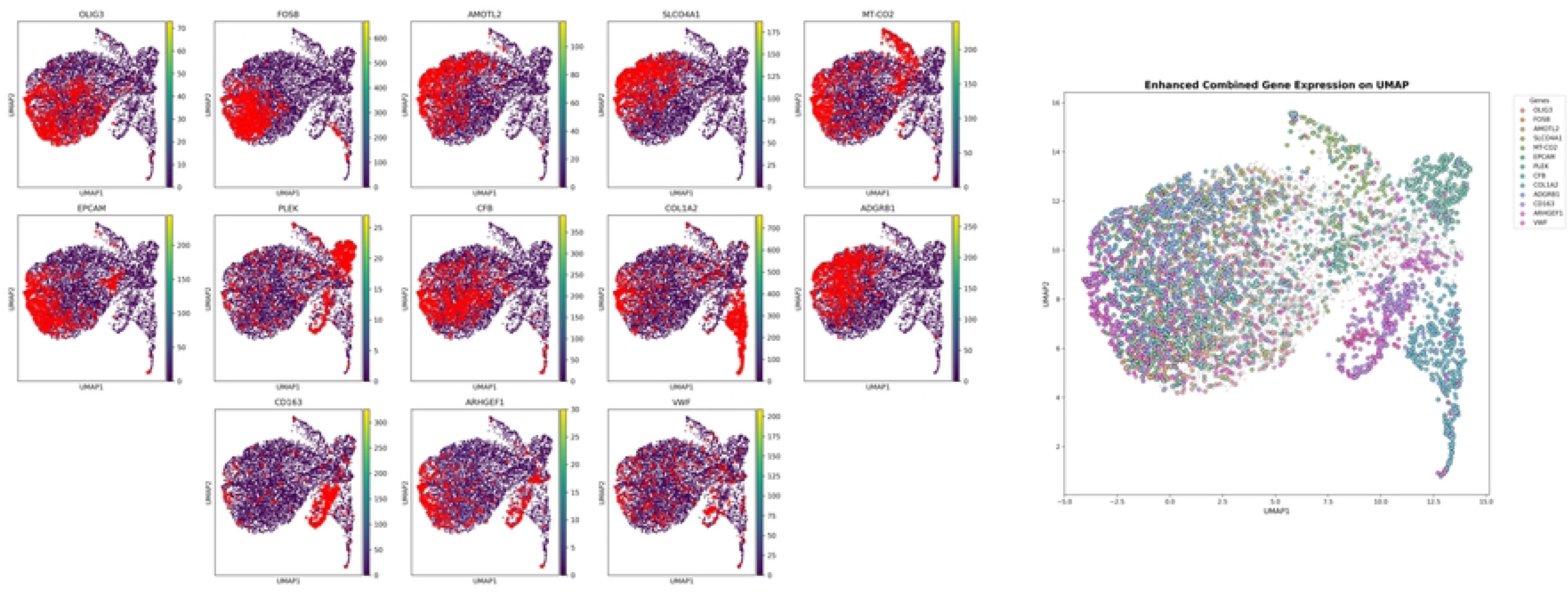
Mapping of High Expression Regions of 13 Genes on UMA.

## 4. Discussion

This study was conducted to investigate the gene expression profiles and functional properties of different cell clusters in the context of ovarian cancer. Cluster analysis was performed with the Leiden algorithm at different resolutions, and 13 clusters were identified at resolution 0.7. It was observed that at lower resolution values, such as 0.4, the clusters were few in number and did not adequately represent biological diversity. On the other hand, at higher resolution values such as 1.0, it was found that the clusters were excessively fragmented and lost their biological meaning; genes were located in excessively fragmented clusters and were insufficient to define common biological processes. Therefore, resolution 0.7 was selected as optimal and was evaluated as a resolution that represents sufficient biological diversity and preserves the meaningful distribution of genes within the clusters [37]. In addition, within the scope of this study, the best genes in 13 cell clusters were determined in the context of ovarian cancer, and the functional enrichment analysis results of these genes were examined in detail. OLIG3, RAB2A, AC007906.2, ZC3HAV1, and CADM1 genes were associated with the POU3F1 transcription factor binding motif in Cluster 0 and were found to be associated with gene regulation in cellular differentiation. In addition, it has been shown in the literature that POU3F1 plays an important role in the transition from epiblast stem cells to neural progenitor cells and activates neural lineage genes while suppressing BMP and Wnt signaling pathways in this process [38, 39]. In addition, specific expression of OLIG3 in V3 neurons adjacent to the p3 domain (at e11.5) also supports its critical role in neural differentiation processes. A study by Fionda et al. showed that POU3F1 is inducible in cancer cells after genotoxic stress and plays a critical role in regulating DNA damage, reactive oxygen species (ROS) management, cellular senescence, and G2 cell cycle arrest [40]. In particular, these regulatory roles in processes such as ROS and DNA damage response highlight their potential implications in differentiation and stress adaptation mechanisms in the context of ovarian cancer. In our study, the association of the binding motif of POU3F1 with OLIG3 supports that this transcription factor may play a critical regulatory role in ovarian cancer cells. These findings suggest that OLIG3 may play an active role in genetic programs regulated by POU3F1 during cellular differentiation [38].

FOSB, GDF15, HMGCR, VTCN1, and TXNIP genes played important roles in progesterone response processes in Cluster 1. FOSB, as one of the prominent genes in Cluster 1, showed a continuous increase during pseudotime and was associated with cellular differentiation processes. In a study by Shahzad et al., FOSB was shown to promote ovarian cancer growth and metastasis together with IL8 under neurological stress conditions [41]. It has been shown that FOSB increases IL8 expression and angiogenesis under the influence of chronic stress, which accelerates tumor growth and promotes metastasis. The role of FOSB in the context of ovarian cancer in our study parallels these findings in the literature and supports the critical role of this gene in cell differentiation and stress adaptation mechanisms. In particular, the continuously increased expression pattern of FOSB and its links to stress-induced tumor growth and metastasis suggest that the gene can be considered as a therapeutic target. AMOTL2, ITGA5, LAMC2, SLCO4A1, and LAMB3 genes contributed to cell adhesion and maintenance of tissue architecture via ECM-Receptor Interaction in Cluster 2 [42]. AMOTL2 was determined as one of the prominent genes in Cluster 2 in our study and was associated with cell differentiation processes. In a study conducted by Tao et al., it was shown that YAP-mediated signaling pathways, including AMOTL2, have an important effect on the senescence of ovarian cancer cells [43]. In this study, it was stated that senescence in ovarian cancer cells is regulated through the S1PR1-PDK1-LATS1/2-YAP pathway and that YAP provides a critical feedback loop in this process. In addition, it was emphasized that YAP increases the sensitivity of cancer cells to chemotherapy and prevents tumor progression. According to the pseudotime analysis results in our study, continuously increasing expression of AMOTL2 was observed; this indicates that the gene has a potential function in differentiation processes and through the YAP signaling pathway. It is noteworthy that the expression level is lower in the early period, but the activity increases as the pseudotime progresses. Although fluctuations are observed at higher values in the late period, the general trend is upward. This regular increase in expression level, especially during pseudotime, supports that it may play a critical role in cell differentiation [42, 44].

SLCO4A1 [45], PADI1, ANGPTL4, SERPINB9 and ITPKC genes are involved in inflammation and metabolic adaptation by being involved in the glucocorticoid receptor pathway in Cluster 3 [46]. SLCO4A1 has been identified as a potential biomarker for predicting platinum resistance status, particularly in ovarian cancer [47]. In this study conducted with machine learning (ML) approaches, it was stated that the expression levels of SLCO4A1 have a critical role in understanding the treatment outcomes and responses to chemotherapy. In this context, the functions of SLCO4A1 in the glucocorticoid receptor pathway are consistent with its role in inflammation and metabolic adaptation processes and are thought to represent an important mechanism in ovarian cancer progression. MT-CO2, MT-CO3, MT-ND4, MT-ATP6 and MT-CYB genes, which are critical in mitochondrial energy metabolism, were associated with mitochondrial inheritance in Cluster 4 [48]. Furthermore, expression of mitochondrial genes such as MT-CO2 is known to play an important role in progression and therapeutic resistance in ovarian cancer through reprogramming of energy metabolism [49]. In the study by Yang et al., the oncogenic roles of GPR176 in ovarian cancer were examined and GPR176 expression was revealed to be involved in biological processes such as focal adhesion, ECM-receptor interaction and cell adhesion molecules by Gene Set Enrichment Analysis [50]. It is suggested that MT-CO2 may play a role in these processes and may provide a critical link between energy metabolism and cellular invasion in particular. EPCAM, CST3, LAMP1, COL4A1 and CD63 genes, which support lysosomal activities and immune response, were found to be linked to the Azurophil Granule Membrane process in Cluster 5 [51]. In addition, studies have shown that EPCAM is highly expressed in 55-75% of ovarian cancer cells and that this gene has the potential to be used in both diagnostic imaging and targeted therapies. Especially in experiments with DARPin Ec1 agent, the high binding specificity and tumor targeting ability of EPCAM are promising for therapeutic applications related to ovarian cancer [52].

PLEK, MXD1, ITGAX, SQSTM1 and PLAUR genes were involved in Protein Kinase C binding processes in Cluster *6* [53]. At the same time, the PLEK gene has come to the forefront with studies on high-grade serous ovarian carcinoma (HGSOC) cell lines related to ovarian cancer. In the study of Masood et al., it was shown that PLEK plays a critical role in cellular signaling and immune responses and can also be evaluated as a potential biomarker and therapeutic target for HGSOC. In particular, PLEK was reported to be one of the hub genes in protein-protein interaction networks and to be associated with NFAT, NF1 and GABP transcription factors in enrichment analyses [54].

CFB, BIRC3, ATP6V1C2, NAMPT and LEFTY1 genes, which are effective in NAD metabolism catalysis, were identified in Cluster 7. In the study by Senent et al., it was stated that the complement system contributes to progression by regulating the tumor microenvironment in ovarian cancer [55]. The study showed that complement activation supports chronic inflammation, creates an immunosuppressive microenvironment and activates cancer-related signaling pathways. It was stated that complement effectors, especially C3a and C5a, and their receptors play critical roles in ovarian cancer progression [56]. COL1A2, COL3A1, COL6A2, COL1A1 and COL6A3 genes, which are effective in the stability of connective tissue, have been associated with the Extracellular Matrix Constituent process in Cluster 8. In Shahan Mamoor’s study, it was reported that COL1A2 showed significantly higher expression in epithelial ovarian cancer (EOC) and HGSOC subtypes and that this gene may be linked to cancer progression or transformation pathways [57]. It has been emphasized that COL1A2 expression differs between normal fallopian tube tissue and ovarian cancer primary tumors and correlates with overall survival of patients.[58] Genes ADGRB1 [59], NID2, CCDC91, LOX and KISS1R, which regulate collagen binding and connective tissue stability, were identified in Cluster 9. The ability of adhesion G protein-coupled receptors (adhesion GPCRs) to regulate extracellular matrix signaling is consistent with a potential role for ADGRB1 in the stability of connective tissue and tumor microenvironment. Furthermore, it has been shown that adhesion GPCRs can be regulated by epigenetic changes such as DNA methylation, and this mechanism has a critical impact on tumor progression and cellular differentiation processes [60].

CD163, SRGN, STAB1, HK3 and LILRB1 genes, which are involved in cholesterol uptake and apoptotic cell clearance through LDL receptor activity, were included in Cluster 10. The study by Reinartz et al. revealed that CD163 was highly expressed in tumor-associated macrophages (TAMs) in malignant ascites associated with ovarian cancer and this was associated with low relapse-free survival rates. The study indicated that cytokines such as IL-6 and IL-10 induced CD163 expression and contributed to tumor progression by increasing immunosuppression in the tumor microenvironment [61].

Similarly, in the study by Heidarpour et al. on serous ovarian tumors, the number of CD163 positive macrophages was found to be positively correlated with the pathological stage, histological grade, and lymphatic metastasis status of the patients [62]. In addition, increased CD163 expression was associated with decreased five-year survival rates of patients, and it was suggested that this gene could be a prognostic marker. These studies support the important roles of CD163 and related genes in ovarian cancer pathogenesis and potential targets for immunotherapies. ARHGEF1, AKNA, ACAP1, TMC8, and IKZF1 genes were involved in the GNA14-p115RhoGEF complex associated with cytoskeletal dynamics and cell migration in Cluster 11. A study by Cheng et al supports this finding. This study analyzed transcriptomes of ovarian cancer cell lines (IGROV1 and IGROV1-CP) and chemotherapy-sensitive and -resistant ovarian cancer tissues using next-generation sequencing technologies. Genes associated with “epithelial-mesenchymal transition” (GO:0001837) and “cell junction assembly and maintenance” (GO:0034330) were found to be enriched in chemotherapy-resistant tissues and IGROV1-CP cells, whereas apoptosis-associated genes were more expressed in chemotherapy-sensitive tissues. Furthermore, genes such as ARHGEF1 have been associated with signaling pathways mediating cytoskeletal rearrangement and cell migration [63]. VWF, EGFL7, SPARC, CD93, and TMEM255B genes, which contribute to endothelial cell functions, vascular structures, and hemostasis mechanisms, were prominent in Cluster 12. In particular, VWF has been shown to play a potential role in age-related ovarian disorders as a biomarker of follicular atresia and ovarian endothelial cells [64]. This finding highlights the influence of vascular structures on ovarian physiology and indicates that VWF should be considered as a target in diagnostic and therapeutic strategies.

Pseudotime analyses of these selected high-scoring genes formed the basis for understanding cellular differentiation processes and better explaining their genetic functions by examining the temporal changes in their expression levels. The expression levels of genes determined for transition cells were examined in detail depending on their resolution values. At the resolution level of 0.4, it was observed that cell clusters were few in number and large in structure. Therefore, gene expression in transition cells exhibited a more homogeneous distribution. While a significant increase in expression was observed especially in OLIG3 and FOSB genes, differences in this resolution were more limited in other genes. Increasing the resolution levels provided a finer separation of cellular subpopulations, which led to the differentiation of gene expression patterns. The higher number of clusters observed at resolution levels of 0.6 and 0.7 allowed a better understanding of the functional diversity among cells. In particular, genes such as MT-CO2 [65] and EPCAM [66] were highly expressed in certain clusters, suggesting that these genes may play a role in critical processes such as cellular metabolism and tissue homeostasis. At resolution 0.8 and 1.0, cells were separated into smaller clusters. This separation allowed transition cells to exhibit more focused expression of specific genes. Expression of genes such as PLEK, CFB, and ARHGEF1 in transition cells became more pronounced at these resolutions. However, some genes (VWF and CD163) began to show differentiated expression profiles at these resolutions.

When the expression profiles of 13 genes were examined during pseudotime, the roles of genes in cellular differentiation processes were revealed in detail. FOSB [67], AMOTL2 [68] and SLCO4A1 genes showed a continuous increase during pseudotime, suggesting that they may be associated with differentiation or stress adaptation processes of cells; it has been suggested that these genes may be active in metastasis and cell motility. In contrast, PLEK, CFB, and ADGRB1 genes showed relatively constant expression levels during pseudotime. This stability-maintaining structure of PLEK suggests that the gene may play a regulatory role in cellular stabilization and signal transduction. CFB, which does not show a significant increase or decrease trend, indicates that it may play a continuous role in the differentiation process. Similarly, the constant expression of ADGRB1 suggests that the gene may play a critical regulatory role in cellular attachment and cell-cell communication. The common features of these genes suggest that they play central roles in maintaining cellular stability and homeostasis. On the other hand, genes such as MT-CO2 and COL1A2 showed a decrease during pseudotime, indicating that they are more active in the early stages and their roles decrease as differentiation progresses.

In the correlation analysis, the positive or negative relationships between genes provide important clues about their roles in common or opposing biological processes. Gene pairs that show a high positive correlation (0.52), such as OLIG3 and AMOTL2, usually function synergistically in the same processes. For example, it is thought that OLIG3 and AMOTL2 may play parallel roles in cellular differentiation or cell motility processes. Similarly, the positive correlation (0.34) of EPCAM and MT-CO2 genes may indicate that they are involved in mechanisms that support each other in metabolic processes and cellular energy production. In contrast, the low correlation between COL1A2 and other collagen genes suggests that connective tissue regulation may occur independently of other cellular processes. This indicates that connective tissue-specific genes may have different regulatory mechanisms [69]. Negative correlations may indicate that genes are involved in opposing biological processes. For example, the negative correlation between VWF and MT-CO2 genes (−0.44) suggests that one is active in vascular structures and hemostasis, while the other plays a critical role in energy metabolism. Similarly, the negative correlation between PLEK and OLIG3 (−0.26) suggests that PLEK may have a fixed role in cellular stabilization or immune responses, while OLIG3 is a more active gene in differentiation or specific cell type development. Such contrasts may reflect that genes exhibit variable expression at different time points or depending on cellular environmental conditions and function in opposing processes [70]. The obtained results provide a valuable resource for better understanding the molecular dynamics of ovarian cancer, while also providing an important basis for the evaluation of these genes as diagnostic, prognostic or therapeutic targets. These findings once again emphasize the importance of multidisciplinary approaches and advanced analytical methods in understanding the biological basis of a complex disease such as ovarian cancer.

## Conclusion

This study aimed to understand the molecular mechanisms underlying processes such as tumor progression, cellular differentiation, and immune response by examining gene expression profiles and functional properties of different cell clusters in the context of ovarian cancer. In the study, Leiden clustering and pseudotime analyses were applied at different resolution levels, revealing biologically meaningful clusters and critical genes associated with cellular processes. It was observed that genes such as FOSB, AMOTL2, and MT-CO2 may play a role in processes such as metastasis, energy metabolism, and cell motility, while genes such as PLEK, CFB, and ADGRB1 may play stable roles in cellular stabilization and signal transduction. The most important advantage of the study is that it provides detailed inferences on both general cellular dynamics and gene basis by integrating clustering, pseudotime analysis, and functional enrichment methods. Resolution-based analyses revealed the biological diversity of cellular populations in more detail and allowed the identification of genes involved in specific processes. In addition, the findings associated with the literature supported the biological and clinical significance of the genes identified in the study and contributed to their evaluation as potential targets. This study has some limitations. Reliance on single cell transcriptomic analyses limits the full understanding of protein-level activity and spatial organization of cells in the tumor microenvironment. In addition, although pseudotime analyses reveal temporal changes of genes, they do not provide direct evidence of functional roles or causal relationships of these genes. The lack of in vitro or in vivo experimental validations in the study limits the applicability of the findings to clinical or experimental contexts. Future studies should aim to validate the roles of genes and pathways identified in this study in in vitro and in vivo models. For example, overexpression or knockdown studies of genes such as FOSB, AMOTL2, or MT-CO2 can be conducted to investigate their functions in tumor progression, metastasis, and stress responses in cell lines. In addition, co-culture systems containing immune and stromal cells may be useful in understanding the interactions of genes, especially CD163 and VWF, with the tumor microenvironment. Spatial transcriptomic and proteomic approaches can provide complementary information on these genes’ cellular localization and protein-level activities. Finally, integrating clinical data with these findings may contribute to the development of personalized medical approaches in ovarian cancer by identifying prognostic and therapeutic biomarkers. This study highlights the value of resolution-based analyses and pseudotime approaches in understanding tumor biology and offers promising avenues for experimental validation and translational research in the fight against ovarian cancer.

## Data Availability

This study’s raw spatial transcriptomics data were obtained from 10x Genomics, specifically the “FFPE Human Ovarian Cancer Data with Human Immuno-Oncology Profiling Panel and Custom Add-on” dataset. This dataset includes comprehensive information on gene expression analyzed using Xenium Onboard Analysis 2.0. The dataset is publicly accessible via the 10x Genomics website. Further, processed data and analysis results generated during this study are available from the corresponding author upon reasonable request.

## Ethics Statement

This study used publicly available spatial transcriptomic data from 10x Genomics’ Xenium platform accessed on 10/08/2024. No direct human participant involvement occurred in our research. The datasets were fully anonymized prior to our access, with no identifying information available to the researchers. As this study focused solely on computational analysis of pre-existing anonymized data, institutional review board approval was not required.

## Code Availability

The code used for data processing, analysis, and visualization in this study is publicly available on GitHub at https://github.com/tissueandcells/OvarySpatial. This repository contains all the necessary scripts and instructions to replicate the analysis presented in this paper. Users are encouraged to cite this work if the code is used in other research.

## Author Contributions

**Conceptualization:** Kevser Kübra Kırboğa.

**Data curation:** Kevser Kübra Kırboğa.

**Methodology:** Kevser Kübra Kırboğa, Emre Aktaş.

**Software:** Ecir Uğur Küçüksille.

**Writing – original draft:** Kevser Kübra Kırboğa, Emre Aktaş, Mithun Rudrapal.

**Writing – review & editing:** Raghu Ram Achar, Gouri Deshpande, Victor Stupin, Ekaterina Silina.

